# A quantitative model of human neurodegenerative diseases involving protein aggregation

**DOI:** 10.1101/541243

**Authors:** Kasper P. Kepp

## Abstract

Human neurodegenerative diseases such as Alzheimer’s disease, Parkinson’s disease, and Amyotrophic lateral sclerosis involve protein aggregation and share many other similarities. It is widely assumed that the protein aggregates exhibit a specific molecular mode of toxic action. This paper presents a simple mathematical model arguing that clinical cognitive status relates to the energy available after subtracting cell maintenance due to general turnover of the misfolded proteins, rather than a specific toxic molecular action *per se*. Proteomic cost minimization can explain why highly expressed proteins changed less during evolution, leaving more energy for reproducing microorganisms on longer evolutionary timescales. In higher organisms, the excess energy instead defines cognitive capability, and the same equations remarkably apply. Proteomic cost minimization can explain why late-onset neurodegenerative diseases involve protein aggregation. The model rationalizes clinical ages of symptom onset for patients carrying pathogenic protein mutations: Unstable or aggregation-prone mutations confer a direct energy cost of turnover, but other risk modifiers also change the available cellular energy as ultimately defining clinical outcome. Proteomic cost minimization is consistent with current views on biomarker histories, explains conflicting data on overexpression models, and is supported by specific experiments showing that proteasome activity is required to confer toxicity to pathogenic mutants. The mechanism and model lend promise to a quantitative personalized medicine of neurodegenerative disease.

## 1. Introduction

Evolution is the fundamental process that has shaped proteomes by randomly mutating DNA and, by random drift or active selection, fixating some mutations in the populations[1,2]. The most prominent relationship of proteomics is that proteins that are highly expressed (i.e. featuring high copy numbers in cells) are more evolutionary conserved in prokaryotes[3] and eukaryotes[4], including mammals[5,6]. This so-called expression-rate (ER) anti-correlation shows that the abundance of proteins in a cell is somehow particularly important during the course of evolution[4,7]. Proteins also tend to use synthetically cheaper (in terms of ATP) amino acids in all kingdoms of life[8,9].

Apparently unrelated to this, millions of people worldwide suffer from age-induced neurodegenerative diseases associated with large deposits of aggregated proteins[10,11]. In Alzheimer’s disease (AD), the so-called senile plaques consist mainly of regular β-sheet fibrils of the infamous β-amyloid peptide (Aβ); in Parkinson’s disease (PD), the Lewis bodies are made up of the protein α-synuclein; in amyotrophic lateral sclerosis (ALS) the deposits consist of superoxide dismutase 1 (SOD1)[12]. Consensus is establishing that these deposits are typically not pathogenic (although toxic) but that soluble protein oligomers inside the cells may attain a pathogenic molecular mode of action. The many histopathological and clinical similarities of these diseases (protein deposits, age-triggering, oxidative stress associated, neurodegenerative) imply a common pathogenesis of protein misfolding[10,13].

Whereas aggregation is cytotoxic and potentially related to disease[14–17], the cellular pathogenic mechanism is unknown and not yet understood[10,14,18–23]. Efforts to identify and target the supposed malicious protein aggregates and oligomers, while possible with inhibitors or antibodies, have so far met with clinical failure[24–29]. This begs the question whether we miss a key determinant of disease course, and it is widely debated whether the protein aggregates in any particular shape and size are the root cause of disease or merely a side effect or contributing feature[10,28–30].

The hallmark protein deposits of the diseases suggest that protein homeostasis (also called proteostasis) is overwhelmed by an excess of misfolded proteins[31]. It is widely assumed that the misfolded proteins are pathogenic themselves by some specific molecular mode of action[32,33]. Although many suggestions have arisen[34], a single, specific molecular mechanism of action has not been identified, and accordingly there are no causative treatments available. Focus now centers on the intracellular oligomers that precede the fibrils; these oligomers are soluble and more toxic to cultured cells than fibrils[34–36]. The oligomers are targeted without actually knowing their bioactive structure, exact location, and the pathogenic process in which they are involved. They interact with many other molecules and cell parts, preventing the identification of a single pathogenic mode of *in vivo* action[27,37]. Efforts to reduce the production of the misfolding proteins by inhibitors or reduce oligomer activity by antibodies, which work in simpler tests, have failed to benefit the human patient[26,28,29,38].

The introduction of a new misfolded protein can overwhelm the existing proteostasis[39]; and some protein misfolding diseases may be transmitted (e.g. prion diseases and controversially suggested, AD [40,41]). Considering the variety of chemical targets within a cell, it is hard to perceive a single molecular mode of action caused by seeding– the mechanism must be generic and involve many kinds of proteins. Such observations beg a formal framework of understanding. The simplest possibility is that oligomers act by a generic mechanism such as membrane permeability disruption by pore formation[42–44], but this still requires a connection to the clinical outcome.

It is widely assumed that the protein aggregation itself is pathogenic, because of the toxicity seen in cultured cells and because genetic mutations causing severe early-onset disease occur in the aggregated proteins (e.g. SOD1 in ALS, α-synuclein in PD, APP and PSEN1 in AD). The mutations, due to their penetration and common dominant inheritance, are often though to gain a toxic function[45–47], although a loss of natural function is also debated for these proteins[30,48–50].

## 2. The model of proteomic cost minimization

The model of proteomic cost minimization as a basis for protein evolution[51] is briefly reviewed below. The E-R anti-correlation has been explained[52,53] as a selection against inefficient translation leading to misfolded proteins. Highly expressed proteins would then be under a stronger selection pressure since the copy number of misfolded proteins *U*_i_ scales with the total abundance of the protein *A*_i_. An empirical equation for the fitness cost *Φ*_i_ has thus been suggested[52]:

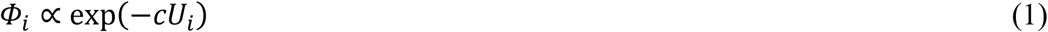

In this equation, *c* is an unknown dimensionless fitness cost constant of one misfolded protein.

To understand this observation at the molecular level, we first introduce a simplified protein turnover scheme, which provides the timeframe of the problem. The proteostasis of any protein *i* can be written in a simplified way as[51]:

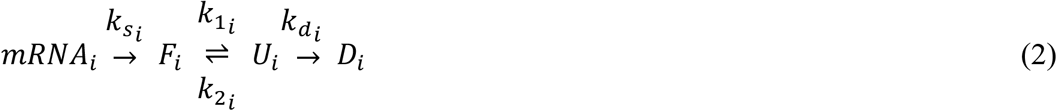

In this scheme, *F*_*i*_ represents the folded protein copies within the cell, *U*_*i*_ represents misfolded protein copies, and *D*_*i*_ are degraded chemical products, with the rate constants of each step annotated. 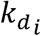 is the rate constant (in units of protein molecules per second) for degradation of the misfolded protein copies. Since many values of degradation rate constants of proteins are known, we also know that the values of 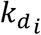 vary greatly with the protein *i*.

We now introduce an estimate of *U*_i_ assuming that the misfolded protein arises from a simple two-state folding process. If so, we have:

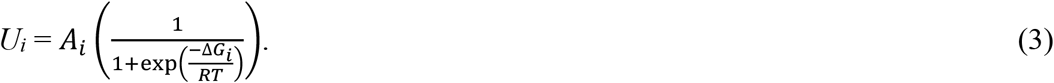

*A*_i_ represents the total copy number of the protein within the cell, Δ*G*_*i*_ is the folding stability (measured as a negative number in kJ/mol) of this protein and *RT* is the thermal energy of the cell at 37°C. In the limit of infinite protein stability (Δ*G*_*i*_ → - ∞), *U*_i_ → 0. The cellular maintenance energy (in J s^-1^) allocated to one protein *i* per time unit can then be derived as[51]:

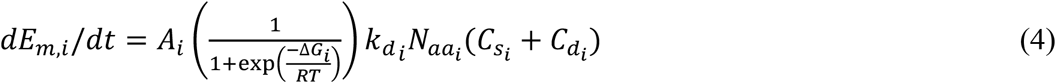

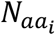 is the number of amino acids in protein *i*, whereas 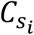 and 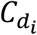 represent the average synthetic and degradation cost (in units of J) per amino acid in protein *i*[51].

Equation (4) is for one protein. Let us now write the *total* protein turnover cost per time unit as the sum of the turnover costs of all proteins within the cell:

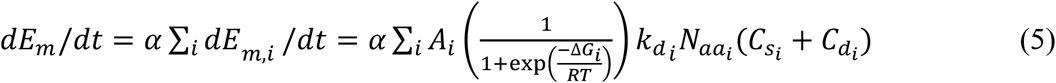

Energy costs are seen to scale with *A*_i_. Copy numbers *A*_i_ vary greatly with the type of protein *i* (typically from zero to 10^6^) and thus some proteins are substantially more systemically important to the energy state as explained by Equation (5). We have included the possibility that the activity of protein turnover affects the total cost by a constant representing an effective concentration of proteases as determining the activity of the proteasome, called *α* in Equation (5). This constant multiplies with 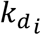 and thus scales the energy cost per time unit.

The energy cost per time unit of the proteome, Equation (5), explains the E-R correlation: One first assumes that the fitness is given by the energy available for reproduction *dE*_*r*_/*dt* after subtracting the energy cost of Equation (5) (ignoring non-proteome energy costs) from the total produced energy of the cell *dE*_*t*_/*dt*:

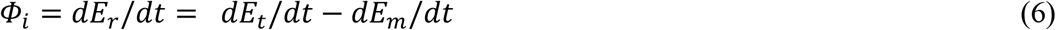

In the simple case where the total energy production is constant, minimization of *dE*_*m*_/*dt* maximizes fitness. From Equation (4), fitness (and hence the selection coefficient) scales with the abundance of the protein. Accordingly, abundant proteins are under stronger selection pressure for cost minimization, making them less likely to accept random (on average destabilizing) mutations and thus conserving them more over evolutionary time. Because the exponential of Equation (1) can be expanded as 1− c*U*_i_ due to the small values of c*U*_i_, this theory recovers and explains the phenomenological fitness cost constant *c* of Equation (1) in terms of fundamental protein turnover parameters[51].

Among costly processes, protein synthesis accounts for ~30% of resting energy expenditure in man[54,55] and typically ~75% in growing microorganisms[56]. Typical costs are ~10 kJ per gram protein, at about 3-5 grams of protein produced per kg mass per day[55]. A typical adult human thus synthesizes 200-300 grams of protein per day, spending 2000-3000 kJ on this process alone per day out of a total basic metabolic rate of perhaps 6000 kJ/day^3^. Protein degradation may cost another 20% of the mammalian total energy expenditure[57], making protein turnover the most energy-consuming process of the body.

## 3. Energy model of neurodegenerative disease

In the following, the fitness function of Equation (6) is argued to also describe a cognitive status function of the brain. Equation (6) defines the energy available after subtracting maintenance costs from the produced energy. Now instead of microorganisms spending their surplus energy on reproduction, higher organisms use a large part (~20%) of their surplus energy for cognition and brain function [58–61]. In higher organisms, we can then consider *dE*_*r*_/*dt* as energy available for cognitive execution, dominated by the ~50% energy spent on the ion pumps[59,62]. To distinguish it, we call it *dE*_*x*_/*dt*:

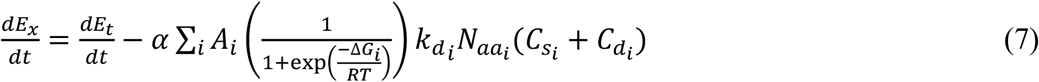

In this simplest model, the energy available for cognition has been reduced to a function only of proteome turnover costs, summed over all proteins *i*. Of course, other costs can easily be included in such a model, but the purpose at this point is to show how protein misfolding relates to the clinical outcome, which we take as *dE*_*x*_/*dt*. Thus, any impairment of *dE*_*x*_/*dt* caused either by a reduction in *dE*_*t*_/*dt* or an increase in maintenance energy *dE*_*m*_/*dt* will reduce cognitive capacity.

From Equation (7), any protein that is associated with an increased abundance *A*_*i*_, turnover constant 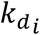, loss of stability to increase the pool of unfolded proteins, Equation (5), or an increase in the specific synthesis and degradation costs of the amino acids 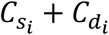 will increase maintenance. Please note that the bias towards synthetically cheap amino acids seen in real protein sequences is largely reflected in the parameter 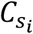 Many other risk factors can be expected to influence either *dE*_*m*_/*dt* or *dE*_*t*_/*dt*. A summary of this mechanism of neurodegeneration is given in **Figure 1**, with energy status being the central determinant of cell fate and cognitive capacity.

**Figure 1.**
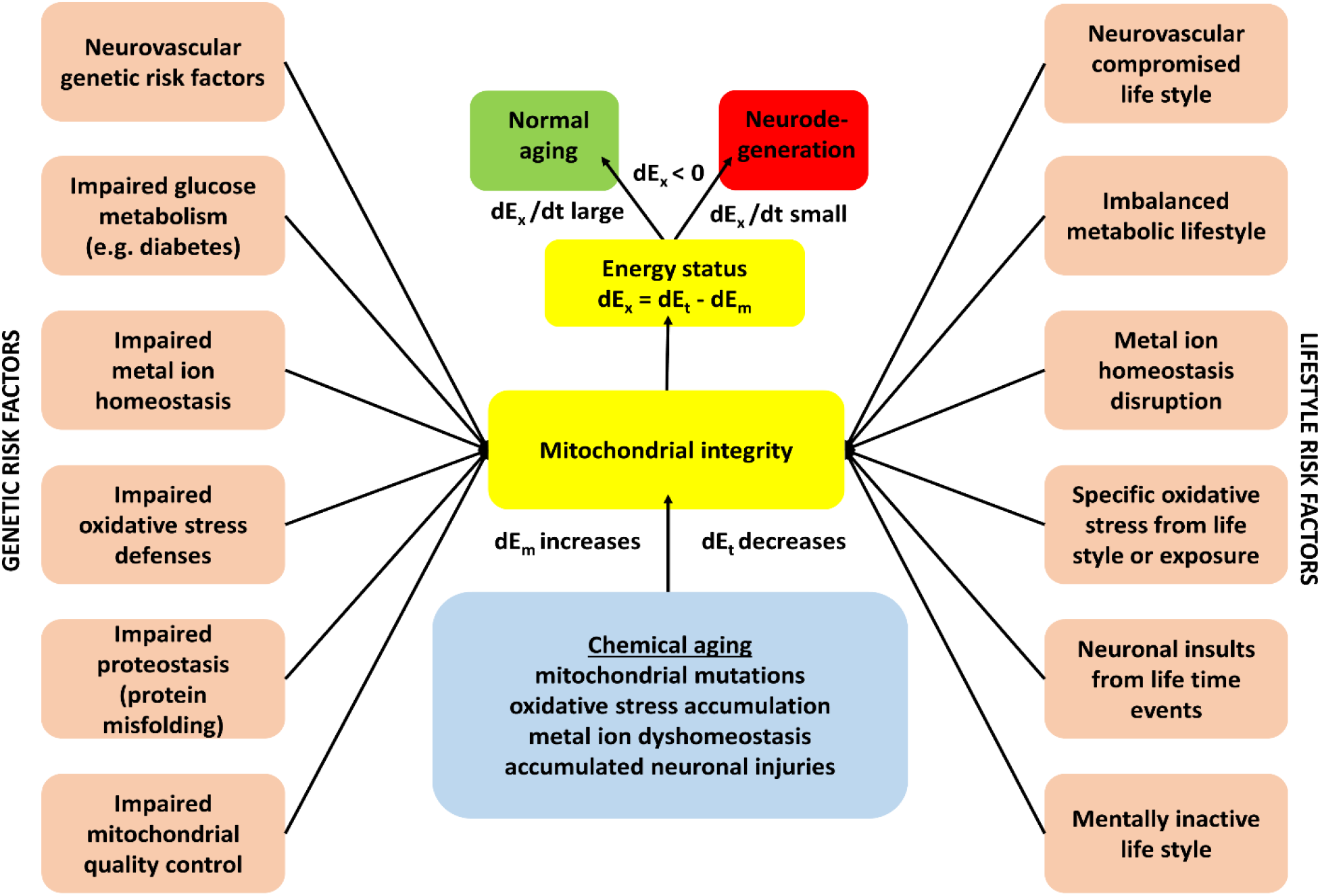
Summary of the exhaustion mechanism of neurodegeneration. Genetic and life style risk factors affect the energy balance of the brain by elevating maintenance costs (dE_m_) or lowering total energy production (dE_t_). If the process occurs gradually, normal aging is observed. If the process is accelerated, the cost of maintenance will increase faster than the ability to compensate energy loss, leading to neurodegeneration when dE_m_ ~ dE_t_.

Energy production typically decreases with age and the energy needed for maintenance increases with age, as seen from the increase in maintenance and stress-related proteins in the aging human brain[63], particular changes in AD[64], and from the glucose biomarkers[65,66]. For simplicity, we first assume that the deceleration in energy production and the acceleration in maintenance costs are constant over time. If so we have:

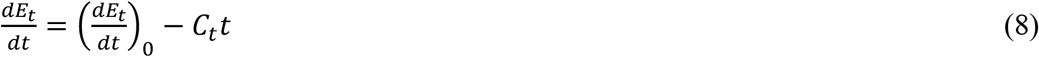

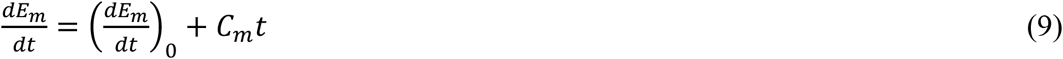

Here, *C*_*t*_ and *C*_*m*_ are the constants of change in the energy production and maintenance costs per time unit, respectively. Accordingly:

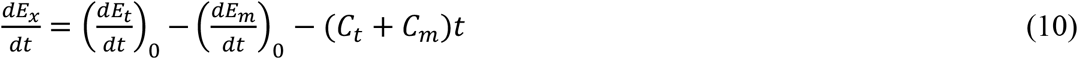

Thus, the energy available for cognition is a function of four variables: The genetically encoded starting respiration rates of energy production and maintenance costs, and the specific constant changes in these two processes:

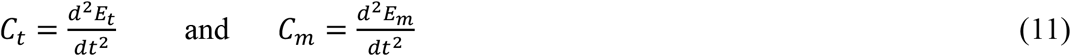

The model is shown in **Figure 2A-C**, using a healthy normality index of 100, with a 50% cost of total energy devoted to maintenance at t = 0. Because of the constant changes in energy production and maintenance, linear changes are seen in d*E*_t_/dt, d*E*_m_/dt, and accordingly, d*E*_x_/dt. For simplicity, the clinical age of onset is assumed to be the point of intersection where d*E*_x_/dt = 0 (dashed lines). The simplest loss of function phenotype would manifest as a larger *C*_t_, the reduction rate of the energy production, whereas the simplest gain of function mechanism would accelerate maintenance costs, *C*_m_. Mixtures of these situations are possible. These two situations correspond to **Figure 2B** and **2C**, where values of *C*_m_ = 0.5 and *C*_t_ = 0.5 have been used respectively, whereas in the “normal” case of **Figure 2A**, *C*_m_ = 0.25 and *C*_t_ = 0.25. The approximate relationship dE_m_/dt ~ dE_x_/dt ~ ½ dE_t_/dt holds for a normal brain[59,62].

**Figure 2.**
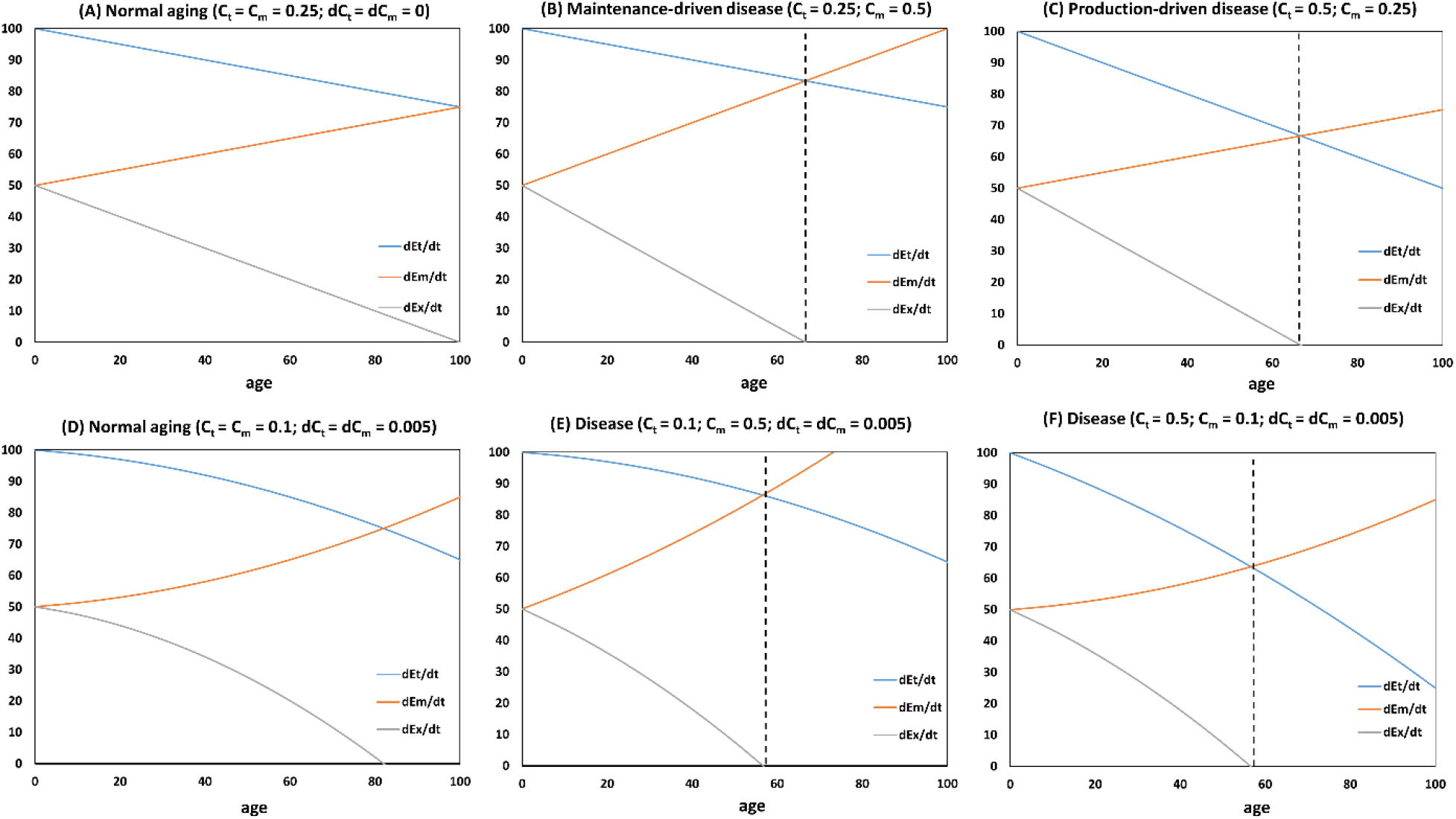
Six scenarios of proteomic cost driven neurodegenerative disease. **(A)** Normal aging with constant changes in energy production and maintenance (*C*_t_ = *C*_m_ = 0.25; d*C*_t_ = d*C*_m_ = 0); **(B)** disease driven by increased maintenance costs (*C*_t_ = 0.25; *C*_m_ = 0.5; d*C*_t_ = d*C*_m_ = 0)**; (C)** disease driven by reduced energy production (*C*_t_ = 0.5; *C*_m_ = 0.25; d*C*_t_ = d*C*_m_ = 0); (**D**) model of accelerated aging (“viscous cycle”) where the rate of change varies (d*C*_t_ = d*C*_m_ = 0.005) for a normal person (*C*_t_ = *C*_m_ = 0.1); (**E**) as in (**D**) but with *C*_m_ = 0.5; (**F**) as in (**D**) but for a patient with *C*_t_ = 0.5.

The age of clinical symptom onset is defined as the time where the surplus energy available for cognition becomes zero, *dE*_*x*_/*dt* = 0:

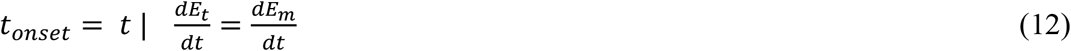

This situation can importantly be modeled as simple constant acceleration as seen in **Figure 2A-2C**. In this simple case, the age of clinical onset simply becomes:

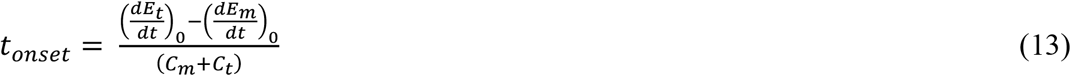

The parameters are the average values for the full cells of the affected cell types. Clinical age of onset depends on the basic energy production and maintenance costs, modified by the accelerated costs and decelerated energy production during aging. All four terms can vary between individual patients as a function of risk modifiers.

Let us for simplicity consider the situation where the patient has normal energy production throughout life, but has increasingly elevated maintenance costs. This can be due to either increased basic maintenance costs or an elevated rate of increase in maintenance costs, the latter shown in **Figure 2B**, described as:

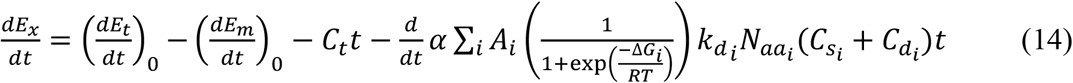

The case in **Figure 2B** represents increased *C*_m_. Several of the parameters in Equation (14) can easily be time-dependent. For example, the amount of misfolded protein may increase due to destabilization caused by oxidative stress other forms of chemical aging. Proteins that are easily and quickly degraded (large 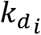) pose a particular challenge to the maintenance costs, but in proportion to their abundance. Thus, the combination of abundant and quickly degraded proteins is the most severe problem in Equation (14).

The model can estimate the clinical age of onset of carriers of familial disease mutations as a function of the severity of a protein mutant. For example, SOD1 mutants causing severe FALS generally exhibit a reduced stability or net charge, which both work to increase the misfolding and turnover of this important and abundant (i.e. large *A*_i_) protein[45,67,68]. The increased cost of maintaining the mutant SOD1 copies folded causes motor neurons to have less energy available for execution over longer time, when total energy budgets become stressed (**Figure 2**). Typical SOD1 mutants reduce Δ*G*_*i*_ by 5-10 kJ/mol[69–71].

A typical decrease Δ*G*_*i*_ of 5 kJ/mol will double the copy number of misfolded proteins in the cell (Equation 3) regardless of the abundance, and in any reasonable range of Δ*G*_*i*_ (−20 to −100 kJ/mol) so this effect is generic. Equation (13) measures the total *C*_m_ for the cell; a doubling of cost of one protein will raise total *C*_m_ by a very small amount. An abundant protein with 100,000 copies would still constitute less than a percentage of total protein in the cell. An increase in total *C*_m_ of 1% would not drive disease in a linear model (**Figure 2A-2C**). A simple linear model is also unlikely considering the inherent accelerations in energy depletion and oligomer buildup: Any deficiency of energy will leave more oligomers and accelerate maintenance cost over time. In such a process, oligomers and associated energy cost build up exponentially until saturation:

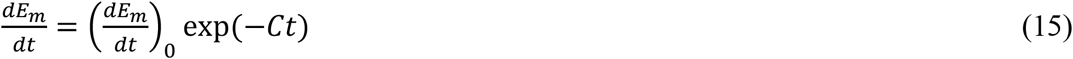

where *C* is the depletion rate constant. Biomarker studies suggest that the disease progression increases nonlinearly and saturates at long times, giving a sigmoidal shape[72].

Let us therefore study a model where *C*_m_ and *C*_t_ increase with time, due to accelerating turnover costs per time. The mechanistic basis is that lack of energy can affect other features such as depolarization of neurons, excitotoxicity, calcium, zinc, and copper imbalances, and oxidative stress that may feed back and aggravate the continuous loss of dE_x_/dt [43,73–76]. **Figure 2D-2F** shows the model where the change in maintenance cost accelerates with aging to produce non-linear, accelerated disease progressions. Even a small acceleration of the maintenance costs can move clinical symptom onset to much earlier age. Mutations that not only increase turnover costs but do so increasingly with chemical aging is described by this situation[77–80]. Whereas the linear scenario is unlikely to lead to disease via a single protein’s impact on the total value of *C*_m_, the non-linear scenario in **Figure 2D-2F** is feasible even at small initial perturbations of the proteostatic machinery.

## 4. Modeling disease histories and biomarkers

The energy model of neurodegeneration as formulated above implies that disease can be more or less severe, depending on the rate of depletion of energy resources. Also, the many risk modifiers will modulate the disease history of a patient in a way that may be modeled. **Figure 3** illustrates a simple disease history for AD, based on the much used framework by Jack et al.[72] Glucose uptake is impaired as the earliest measurable event, followed by Aβ pathology[72]. A patient with a mutation that produces oligomers subject to intensive proteolysis will exhaust the energy status of the neurons faster than the typical patient; this would lead to an early-onset form of the disease, shifting the age of onset towards the left of **Figure 3**. Since this clinical age of onset is directly defined by Equation 12, using for illustration an indexed level of function loss of 0.15 as threshold where disease symptoms are evident. In this way, the modeled disease progressions in **Figure 2** can be mapped directly to the clinical disease progression of **Figure 3**.

**Figure 3.**
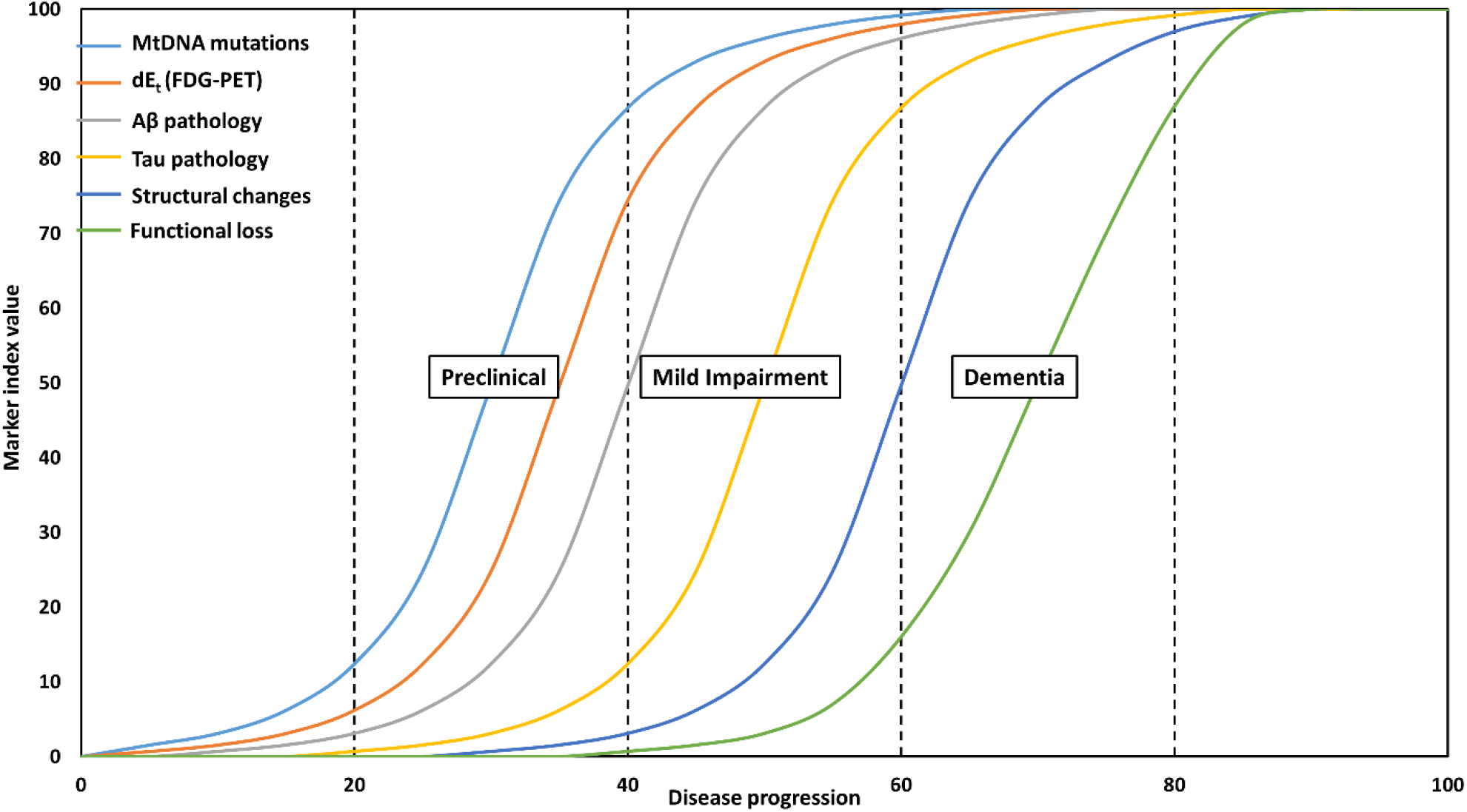
Approximate biomarker normalized index value (0-100) vs. hypothetical disease progression (from 0 to 100), based on a previous model[72]. Age of onset is arbitrarily chosen to be the point at which indexed functional loss is 0.15 (marked in stipulated lines as separating mild impairment from dementia) and can be estimated from Equation (13)-(15).

To understand this in more detail, the non-linear progression of biomarkers[72] follows directly from the exponential aging model represented by the accelerated proteomic energy cost, **Figure 2E** and **2F**: At any age, the buildup of pathogenic protein oligomers will accelerate because the energy left for degrading them decreases exponentially (**Figure 2E, 2F**). Because these oligomers are in an equilibrium with aggregated deposits, the deposits will increase exponentially until saturation is reached when the amount of aggregated protein does not increase further because cell death cancels the new production of aggregates and partly because the export of aggregates becomes unfavorable as the extracellular aggregate concentration increases. Thus the model rationalizes the clinical observed history of biomarkers.

The common explanation given for the sigmoidal shape of biomarkers [72] is that the shape parallels the *in vitro* aggregation assays, typically monitored by the β-sheet binding thioflavin T, such that an elongation phase is followed by nucleation, leading to an exponential increase is aggregation until saturation. However, in a real neuron, it is very unlikely that this *in vitro* mechanism takes place considering the intracellular concentrations of proteins that will probably only form large aggregates outside the cell, but be constantly degraded as oligomers inside the cell. The model in **Figure 2** provides an alternative, systemic explanation for the sigmoidal shape not due to aggregation kinetics but to accelerating exhaustion of turnover capacity. These two explanations for the observed sigmoidal biomarker histories are fundamentally different and may be tested in the future.

## 5. Support for the model: Energy in the healthy brain and in disease

The model immediately answers the haunting question why neurodegenerative diseases are protein misfolding diseases. A pool of misfolded proteins always exists in all cells but is kept very small, probably below one copy at any instance per cell due to fast turnover. The energy burden and thus disease only manifests in the most energy-requiring cells, including those of the central nervous system. The adult human brain typically uses 20% of the total resting energy, despite weighing only 1200-1500 grams, a few percent of total body mass[58,81–83]. Typically 50% of this brain energy is allocated to maintain the ion gradients central to neuronal signaling, whereas house-keeping takes up the second-largest contribution and includes protein synthesis and degradation costs[59,62]. The neuronal maintenance costs per time unit set the limit on the cognitive abilities via the temporal efficiency of information processing[58,62]. Despite the very high use of energy, very little energy is stored in the brain, and the neurons are thus extremely sensitive to the supply of energy[84]. Upon circulatory arrest, brain energy stores are used up within a few minutes[60]; thus dE/dt rather than absolute energy stores is used in the model.

Age-induced impaired energy balance is associated with all major neurodegenerative diseases: Glucose abnormalities feature in both PD[65,85] and AD[86–88], and metabolic changes are associated with PD[89], AD[64,90], and ALS[91]. Diabetes and obesity are risk factors of neurodegenerative disease[87,90,92], as are genes related to cholesterol and lipoprotein homeostasis[93,94]. Energy deficiency has been suggested to be a major underlying cause of AD[30,86,95–97] and ALS[98–101], and deficiencies in mitochondrial respiration is a well-known central feature of PD[89,102–104]. Reduced energy availability in AD is well-established and sets in before clinical symptoms[64,77,105].

AD has been viewed by some as a mitochondrial disease[78,106–109], and is strongly related to calcium dyshomeostasis[110–114]. Abnormal energy availability in the brain is a consistent feature of AD[93,115–117], and potential blood markers of AD relate to apoptosis and metabolism; clusterin, which is a risk factor and candidate biomarker, probably plays a central role in these pathways[118]. AD patients have reduced availability of glucose/insulin[86,97], and disruptions in metabolism and energy balance of the brain is an early feature of AD[90]. The energy shortage is a key feature of AD[64,119], and AD is also characteristically a mitochondrial disease that reduces dE_t_/d_t_ relative to dE_m_/dt [107,120,121].

Combined Aβ and glucose exposure to microvascular endothelial cells cause enhanced cooperative chemical aging[122]. This suggests that Aβ plays a role in glucose metabolism, and Aβ can increase glucose tolerance[123]. One of the strongest lines of evidence for a direct connection between protein misfolding toxicity and accelerated aging related to disease comes from studies of Aβ effects on insulin signalling[124].

PD is most likely caused by impaired ability to quality-control mitochondria of the affected cells, resulting in low energy production and associated energy shortage[89][65]. As is the case of AD and ALS, protein quality control is also impaired in PD[21], as is specifically the quality control of the energy-producing mitochondria, and parkin affects both mitochondrial quality control and glucose metabolism of the brain[85].

SOD1 aggregates in sporadic ALS, and mutations cause severe early-onset familial ALS (FALS)[101,125,126]. ALS patients experience increased resting energy expenditure that has not been rationalized[127],[91], but fits Equation (14) since SOD1 mutants almost invariably decrease stability and thus increase the pool of misfolding proteins requiring turnover. The observations that ALS-causing SOD1 mutants impair mitochondrial respiration[128,129] and cause metabolic abnormalities[130] support the direct relationship between protein turnover of aggregation-prone mutants and cellular energy claimed by Equation (14). Thus, the model suggests that ALS results from degeneration of motor neurons caused by depletion of the very large energy required for executing motor-neuron signaling[131]. There is now substantial support for this explanation[132–134]. Additional elevated costs of turnover will contribute to Equation (14) as well via the proteasome parameter α. As an example of the effect of the latter, functionally impaired variants of Ubiquilin-2 have been implicated as a risk factor in ALS[135].

Any increased turnover of a protein also involves a cost associated with RNA processing. Some mutations do not result in amino acid changes in the produced proteins but still cause disease. The GGGGCC hexanucleotide repeat expansion on chromosome 9, C9ORF72, accounts for many cases of ALS. Such repeat expansions will lead to abnormal RNA processing and transcriptional inefficiency[136], which we argue manifest as an increased ATP cost of the cell per time unit, i.e. an increase in *C*_m_ of Equation (14). Other recently identified genetic risk factors also affect RNA metabolism (TAR-DNA binding protein 43[137,138], FUS[139,140]) or protein processing (SQSTM1[141], VCP[142]). Also, TDP-43, a risk factor in ALS, has been suggested to modulate SOD1 levels[143].

If the model is correct, overexpression of any protein, if substantial enough, should potentially cause energy crisis and be toxic. This relates to the many models of disease where proteins are heavily overexpressed. Overexpression of non-pathogenic SOD1 wild-type proteins produce disease-like phenotypes, although these effects are smaller than for the pathogenic mutants[144]. These observations do not fit well with the standard models presuming an innocent wild type and a pathogenic mutant regardless of expression level, but are completely explained by Equation (14), where both the abundance *A*_i_ and the stability change contribute to the pool of misfolded proteins subject to turnover, and thus to the energy costs.

A simple, central experiment can further test whether the proteomic cost minimization model or the “molecular toxic action” mechanism is correct. It involves expressing pathogenic mutants with and without the protein being degraded. If the proteasome is inhibited during expression of the aggregation-prone mutants, disease would be aggravated if the misfolded proteins were toxic *per se*, because more toxic misfolded proteins would be available. However, if disease is mainly due to the energy burden of turnover, short-term proteasome inhibition concurrent with overexpression of pathogenic mutants should be less pathogenic than the same mutants expressed at the same levels without proteasome inhibition on a short time scale until the inhibition of protein turnover becomes systemically toxic.

It was recently shown that the pathogenic G85R mutant of SOD1 is not very toxic under normal conditions and during inhibition of the proteasome, but when the proteasome activity is recovered after inhibition, the soluble mutant oligomers become toxic[133]. The general assumption that a molecular mode of action makes misfolded proteins toxic is not supported by such experiments. Equation (14) explains it because the toxic mode of action is not by molecular activity of the misfolded protein copy, but by its associated turnover cost. Proteasome inhibition can lead to several-fold higher levels of SOD1 within the cell[145], which shows that the turnover an its disruption readily and to a sufficient extend changes the energy costs.

## 6. Concluding remarks

There is a pressing need for new fundamental understandings of neurodegenerative diseases and for models that can combine and rationalize the vast amount of data in the field. This paper provides such a model relating protein turnover directly to clinical outcome. The principle of proteome cost minimization holds that neurons are among the most energy-demanding cells in the body, that the cost of protein turnover reduces the energy left for ion pumping and thus cell signaling, and that the increased energy costs and reduced energy production produce a threshold where energy available for cognitive execution becomes critical, the age of symptom onset. Abundant, aggregation-prone proteins will increase maintenance cost proportionally and thus accelerate this age. Turning on and off the proteasome by inhibitors during pathogenic mutant expression in cultured cells should be a straightforward way to test the theory.

The equations suggests multiple strategies to address the diseases, notably by either increasing total energy availability, energy efficiency, or by reducing maintenance costs. Therapies that target those oligomers that are particularly subject to proteolysis *in vivo* are predicted to be most advantageous. The predictions are in contrast to those of the traditional view that protein aggregates have a specific toxic mode of action that should be targeted, and that any turnover of them, however costly, should be less important to disease course. Such clinical strategies have so far failed perhaps for the reasons stated in this work. Instead, the increasing evidence for the case of targeting either energy production, availability, or consumption[146] is largely consistent with the model described in the present work.

## Acknowledgements

The author acknowledges financial support from the Novo Nordisk Foundation, grant NNF17OC0028860, and the Danish Council for Independent Research | Natural Sciences (DFF), grant 7016-00079B.

